# CASSIA: a multi-agent large language model for reference free, interpretable, and automated cell annotation of single-cell RNA-sequencing data

**DOI:** 10.1101/2024.12.04.626476

**Authors:** Elliot Xie, Lingxin Cheng, Jack Shireman, Yujia Cai, Jihua Liu, Chitrasen Mohanty, Mahua Dey, Christina Kendziorski

**Affiliations:** Department of Biostatistics and Medical Informatics, University of Wisconsin-Madison, Madison, WI, USA; Department of Neurological Surgery, University of Wisconsin-Madison, Madison, WI, USA

## Abstract

Cell type annotation is an essential step in single-cell RNA-sequencing analysis, and numerous annotation methods are available. Most require a combination of computational and domain-specific expertise, and they frequently yield inconsistent results that can be challenging to interpret. Large language models have the potential to expand accessibility while reducing manual input and improving accuracy, but existing approaches suffer from hyperconfidence, hallucinations, and lack of reasoning. To address these limitations, we developed CASSIA for automated, accurate, and interpretable cell annotation of single-cell RNA-sequencing data. As demonstrated in analyses of over 970 cell types, CASSIA improves annotation accuracy in benchmark datasets as well as complex and rare cell populations, and also provides users with reasoning and quality assessment to ensure interpretability, guard against hallucinations, and calibrate confidence.

## Introduction

Accurate cell type annotation is essential for understanding the cellular composition of tissues, characterizing the functional states of distinct cell types, and identifying dynamic changes in response to disease. Reference-based annotation methods typically rely on existing single-cell RNA-seq datasets with known cell-type labels. Among these, classifier-based methods train supervised machine learning models such as random forests^1^, support vector machines^2^, or deep learning models like transformer-based architectures^3,4^ on labeled reference datasets to classify unknown query cells. Alternatively, correlation-based methods annotate query cells by comparing their expression profiles to labeled reference data without explicit training^5–7^. Reference-based approaches generally perform well given representative reference datasets having features and cell subpopulations that closely match the query data. However, the performance of reference-based methods can be compromised when suitable reference datasets are not available^8,9^.

Reference-free approaches have been developed that leverage prior knowledge of marker genes, thus avoiding reliance on matched reference datasets^10–12^. Whether reference-based or reference-free methods are initially employed, manual annotation often follows to refine those clusters having no or low-confidence annotations, and/or those showing discrepant results across methods^9,13^. While most investigators consider manual annotation by an expert to be the gold standard, it is time consuming, labor intensive, and requires a combination of computational and domain-specific expertise^13^. Large language models (LLM) address a number of these challenges and are being used to streamline and improve computational analyses of genomic data in many domains^14–16^, including cell annotation^17^. Most of these LLMs are built on transformer-based neural networks trained on vast text corpora including scientific literature. Unlike domain-specific models trained on gene expression profiles^3,4^, general-purpose LLMs benefit from continuous improvement through intense competition among leading technology companies. They also possess sophisticated reasoning capabilities that approximate human expert-level analysis, making them particularly valuable for repetitive biological tasks. The popular LLM-based approach GPTCelltype, for example, demonstrates the potential of LLM-based models to reduce the effort and expertise required for cell type annotation^18^. Indeed, with a single-prompt, Hou and Ji demonstrate that GPTCelltype provides annotations in “strong concordance with manual annotations”. In spite of the advantages brought forth by simplifying cell annotation analysis, GPTCelltype has several limitations. A major one is its potential to hallucinate, confidently providing an incorrect annotation. In addition, the approach does not explain the reasoning underlying any given annotation, nor does it provide quality scores. As a result, it is often not possible for a user to distinguish hallucinations or low-quality annotations from high-quality ones. Finally, in an effort to reduce the likelihood of hallucinations, GPTCelltype uses relatively few markers which may be sufficient for annotating general cell types but does not allow for accurate annotation of distinct subpopulations. See Supplementary Note 1 for details.

To address these limitations, we developed a <u>c</u>ollective <u>a</u>gent <u>s</u>ystem for <u>s</u>ingle-cell interpretable <u>a</u>nnotation (CASSIA). In short, CASSIA is a multi-agent LLM framework consisting of five interconnected LLMs for annotation, validation, formatting, quality scoring, and reporting. For applications requiring highly detailed annotations, CASSIA also includes a retrieval-augmented generation (RAG) agent that retrieves external information. Additionally, for cases exhibiting signs of low-quality annotations, CASSIA provides an advanced annotation-boosting agent to further refine and correct results. Optional LLMs are also available for subclustering and uncertainty quantification.

Benchmark analyses in over 970 cell populations show that CASSIA increases cell annotation accuracy, improves interpretability, and provides for robust quality assessment of annotations. In addition to improving the accuracy and ease with which cell annotation can be performed, the utility of CASSIA is further demonstrated in analyses of complex datasets, rare cell types, and mixed cell populations. CASSIA is also used here to identify errors in gold-standard datasets, demonstrating its potential for refining existing reference annotations.

## Results

### Overview of CASSIA

CASSIA is a modular, multi-agent LLM framework for accurate, interpretable, and adaptable cell type annotation in scRNA-seq data. The framework requires a user to specify species, tissue type, groups of cells (cell types) to be annotated, and a collection of markers associated with each cell type, if known. We expect that most analyses will use markers from *FindAllMarkers* in Seurat or *tl.rank_genes_groups* in Scanpy, although markers from other sources may be used if desired. The default CASSIA workflow consists of four main steps:

1. Annotation: Analyze marker expression patterns to generate cell type labels with detailed reasoning.
2. Validation: Iteratively check annotations for consistency in marker-cell type alignment (up to three refinement cycles).
3. Quality Assessment: Assign quality scores (0-100) based on scientific accuracy and marker balance.
4. Refinement: Flag low-scoring or mixed clusters for additional refinement.
5. Reporting: Provide full interpretability with detailed reasoning, quality scores, and refinements.

A schematic illustrating this multi-agent LLM system and workflow is provided in Figure 1. Users provide input information such as species, tissue type, and markers (Figure 1a), which is then processed to create input for annotation, validation, formatting, scoring, and reporting (Figure 1b), culminating in comprehensive output (Figure 1c). For cell types requiring additional refinement, CASSIA offers optional specialized agents (Figure 1d). Specifically, the Annotation Boost agent leverages complete differential expression results such as p-values and fold-changes to improve low-scoring annotations; and the Subclustering agent resolves mixed cell populations with subtle phenotypic differences. For novel or complex datasets where markers are mentioned in the literature relatively infrequently, and therefore are not well represented in GPT and similar LLMs, users can replace the default Annotation agent with a retrieval-augmented generation (RAG) agent that integrates external knowledge from databases like CellMarker^19^ and ontologies^20^, while maintaining the same validation, quality assessment, and reporting workflow.

**Figure 1:**
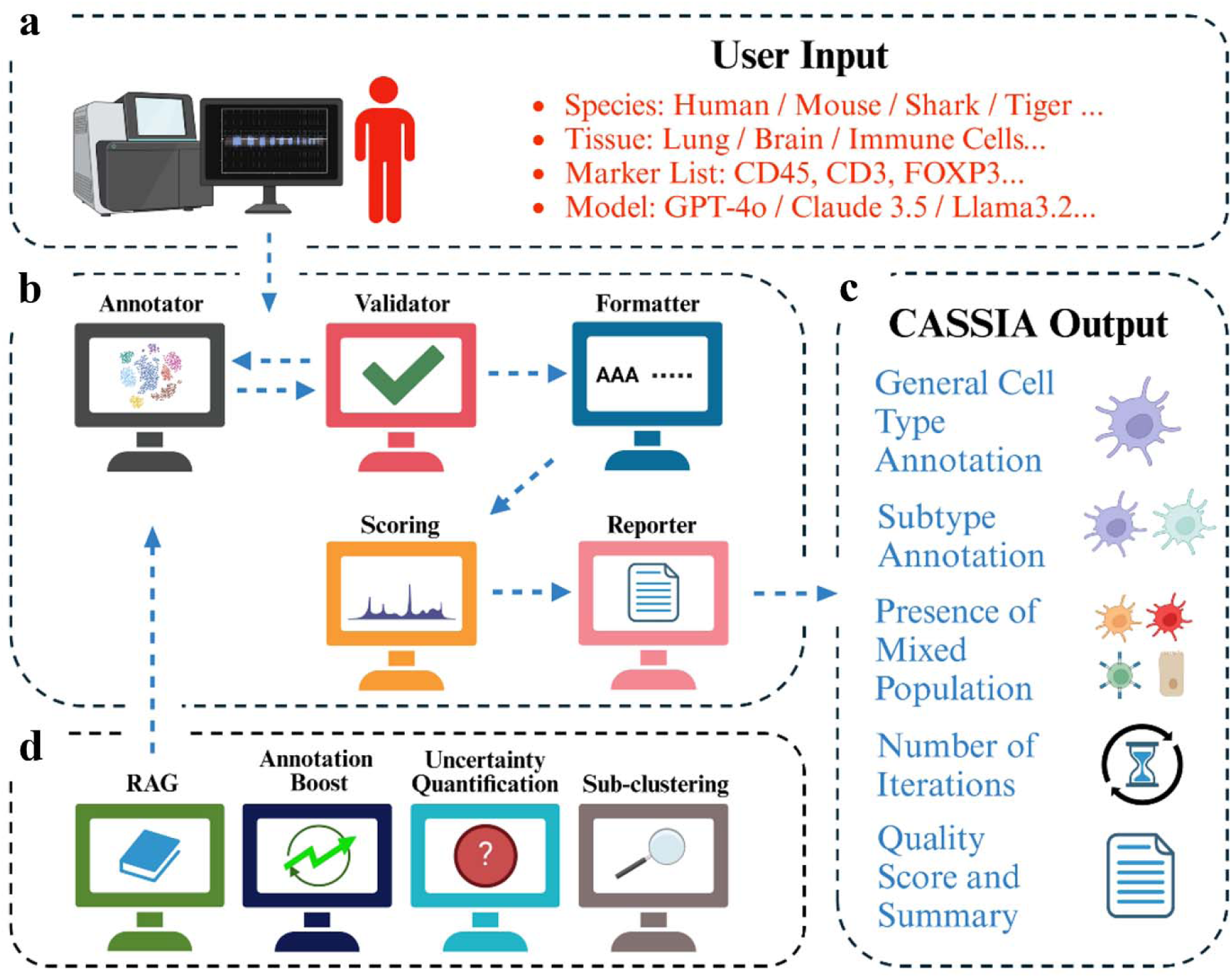
The multi-agent LLM system underlying CASSIA. **(a)** A user interacts directly with the Onboarding platform by specifying species, tissue type, and a collection of markers associated with cell subtypes within that tissue, if known. Any information associated with experimental conditions, interventions, or other sample specific information may be provided. **(b)** Together, this input is used to create the user prompt given to the Annotator agent. The Annotator agent performs a comprehensive annotation of the single-cell data using a zero-shot chain-of-thought approach that mimics the standard workflow that a computational biologist would typically follow for cell annotation. Results are then passed to the Validator agent to check marker and cell type consistency; results failing validation are passed back to the Annotator and this iterative process continues until results pass validation (or the maximum number of iterations is reached). Results are then moved to the Formatting agent which summarizes each cell annotation; this summary along with the full conversation history is provided to the Scoring agent for quality scoring. The Reporter agent then generates a comprehensive report documenting the complete annotation process, including agent conversations, quality evaluation reasoning, and validation decisions with supporting evidence to facilitate transparent interpretation of results. The output from default CASSIA is shown in **(c)**. Optional agents include those shown in **(d)**.

### CASSIA improves annotation accuracy in benchmark datasets

CASSIA was compared with state-of-the-art cell annotation methods on five classic benchmark datasets for which gold-standard annotations are available: GTEx, Tabula Sapiens (TS), Human Cell Landscape (HCL), Mouse Cell Atlas (MCA), and Azimuth. To assess annotation accuracy, we established a hierarchical evaluation framework based on the Cell Ontology tree structure classifying annotations as fully correct, partially correct, or incorrect based on their taxonomic distance from reference annotations. A fully correct annotation matches the reference exactly while an annotation one step away from the reference on the ontology tree is considered partially correct (e.g., predicting “T-cell” for a CD8+ T-cell); annotations more than one step away are classified as incorrect (e.g., predicting “immune cell” for a CD8+ T-cell). See Methods. Figure 2a and 2b show that CASSIA increases fully correct annotations by 12-41% relative to existing approaches and combined correct annotations (fully or partially correct) by 9-29% relative to the next-best approach across benchmark datasets. Performance averaged across all annotations shows that CASSIA increases annotation accuracy by over 20% for most datasets (Figure 2c).

**Figure 2:**
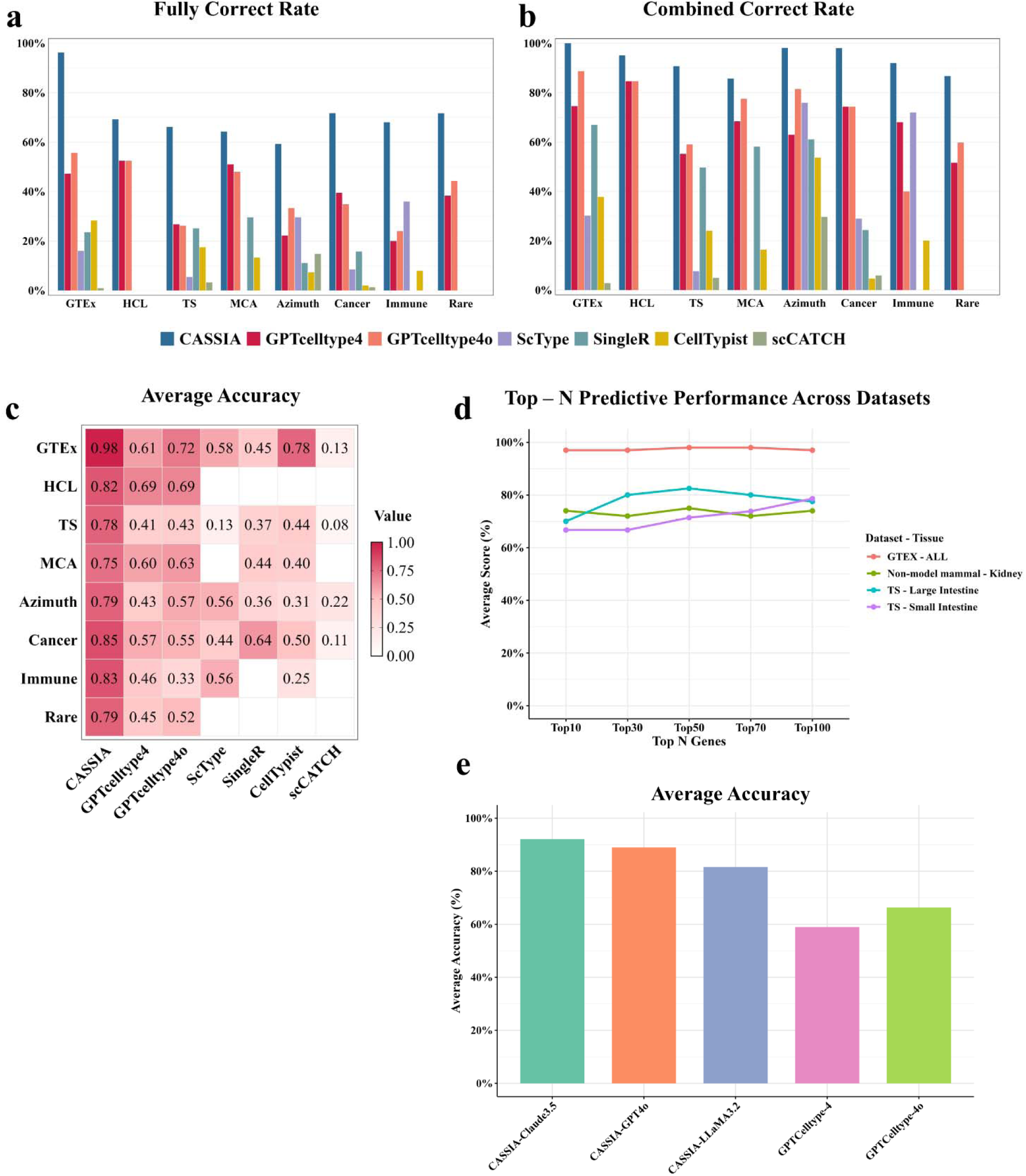
CASSIA improves annotation accuracy in five benchmark datasets, on complex populations of cells from immune and cancer cell populations, and on rare cell types. **(a)** Fully correct annotation rates across 8 datasets where CASSIA (blue) increases annotation accuracy by 12-41% over the next best performing method. **(b)** Combined correct annotation rates (fully or partially correct annotations) across the same datasets where CASSIA maintains a 9-29% advantage over the next-best approach (typically GPTCelltype4/4o). **(c)** Heatmap of average accuracy across cell types for each dataset (rows) and method (columns), with values ranging from 0 (white) to 1 (dark red). CASSIA consistently achieves the highest accuracy scores. **(d)** Top-N predictive performance showing CASSIA’s accuracy across varying numbers of marker genes (10-100) in four datasets. CASSIA maintains consistently high performance (∼0.98) on the GTEx-ALL dataset (red), with stable performance on other tissues. **(e)** Performance benchmarking of CASSIA with different LLMs versus GPTCellType implementations on 100 cell types sampled from five benchmark datasets. CASSIA-Claude-3.5 achieves superior performance (0.92), followed by CASSIA-GPT4.0 (0.88) and CASSIA-LLAMA-3.2 (0.82), demonstrating the framework’s robustness to underlying model selection.

We systematically evaluated the optimal number of marker genes and determined that using 50 marker genes per cell type provides the best balance of accuracy and efficiency. When analyzing clusters with complex cellular compositions or those requiring detailed annotation, it may be useful to include more than 50 markers. Figure 2d demonstrates that CASSIA maintains robust performance even with larger marker sets (e.g., top 100 markers), while competing methods such as GPTCelltype show significant performance degradation due to context window limitations (Supplementary Note 1a). CASSIA’s performance is also robust across different underlying language models, including Claude 3.5 Sonnet, GPT-4o, and open-source LLaMA-3.2 90B (Figure 2e).

Beyond accuracy and robustness, CASSIA provides interpretable results through structured analysis reports that document CASSIA’s reasoning process from functional marker evaluation through cell type marker identification and database validation. Figure 3 shows CASSIA analyzing a colorectal cancer cluster, identifying it as intestinal epithelial cells with a predominant colorectal cancer phenotype. Each annotation includes validation steps with marker consistency checks and pathway verification, along with a confidence score. This transparent process allows researchers to understand the basis for annotations and assess their reliability. Having established a strong baseline in standard benchmark datasets, we next investigated CASSIA’s capabilities in more challenging biological contexts.

**Figure 3:**
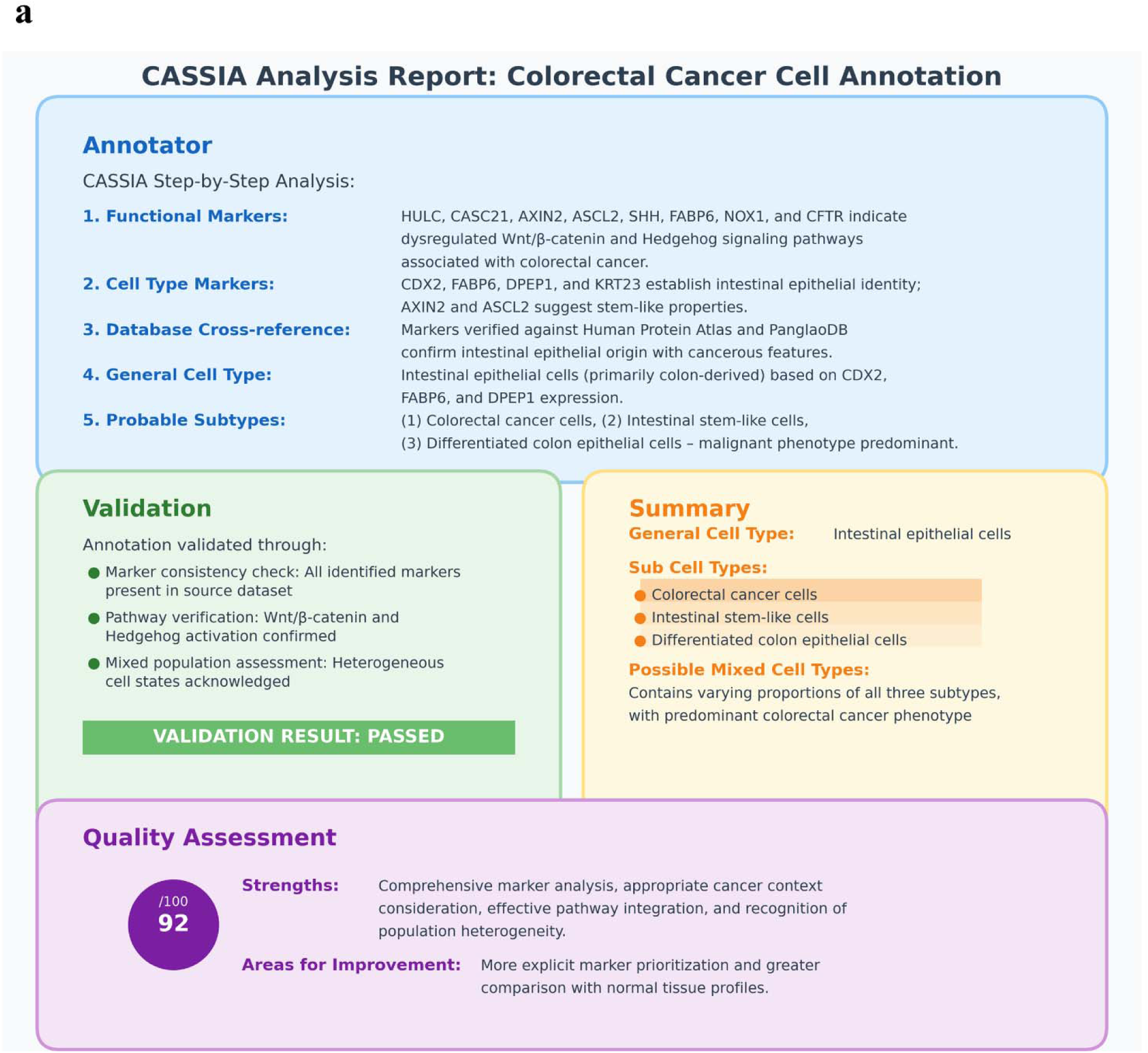
CASSIA analysis report for a cell cluster from a colorectal cancer dataset. The report presents a comprehensive analysis including functional and cell type marker identification, database cross-referencing, cell type determination, and subtype classification. The validation section confirms marker consistency and pathway verification, while the quality assessment provides a numerical score (92/100) with identified strengths and areas for improvement.

### CASSIA performance is maintained in more specialized applications

To systematically assess CASSIA’s performance in more specialized biological contexts, we considered datasets from cancer biology, immunology, and non-model organisms. We first focused on evaluating CASSIA’s performance in cancer datasets composed of two primary cancer samples and three metastatic samples. CASSIA outperforms other methods in assigning correct cell types, achieving 85% accuracy across all cancer datasets (Figure 2c). When tasked with specifically distinguishing cancer from non-cancer cells, CASSIA correctly identifies 72.5% of cancer cells on average across multiple datasets, compared with 20% for GPTCelltype (Figure 4a and Figure 4b). A simple enhancement to CASSIA - adding a single prompt instruction that reads “You should carefully distinguish between cancer cells and normal cells with reasoning” - further improves detection accuracy to 86-100% across all cancer datasets (Figure 4b and Supplementary Data 1). Beyond cell type identification, we also investigated CASSIA’s ability to detect metastasis signals in three datasets. Each consists of single-cell RNA-seq data profiled from metastatic brain tumors with primaries in lung. CASSIA successfully identifies both the metastatic nature of the cells and the tissue of origin in all three datasets (Supplementary Data 1) whereas GPTCelltype accurately detected metastatic cells in only one sample.

**Figure 4:**
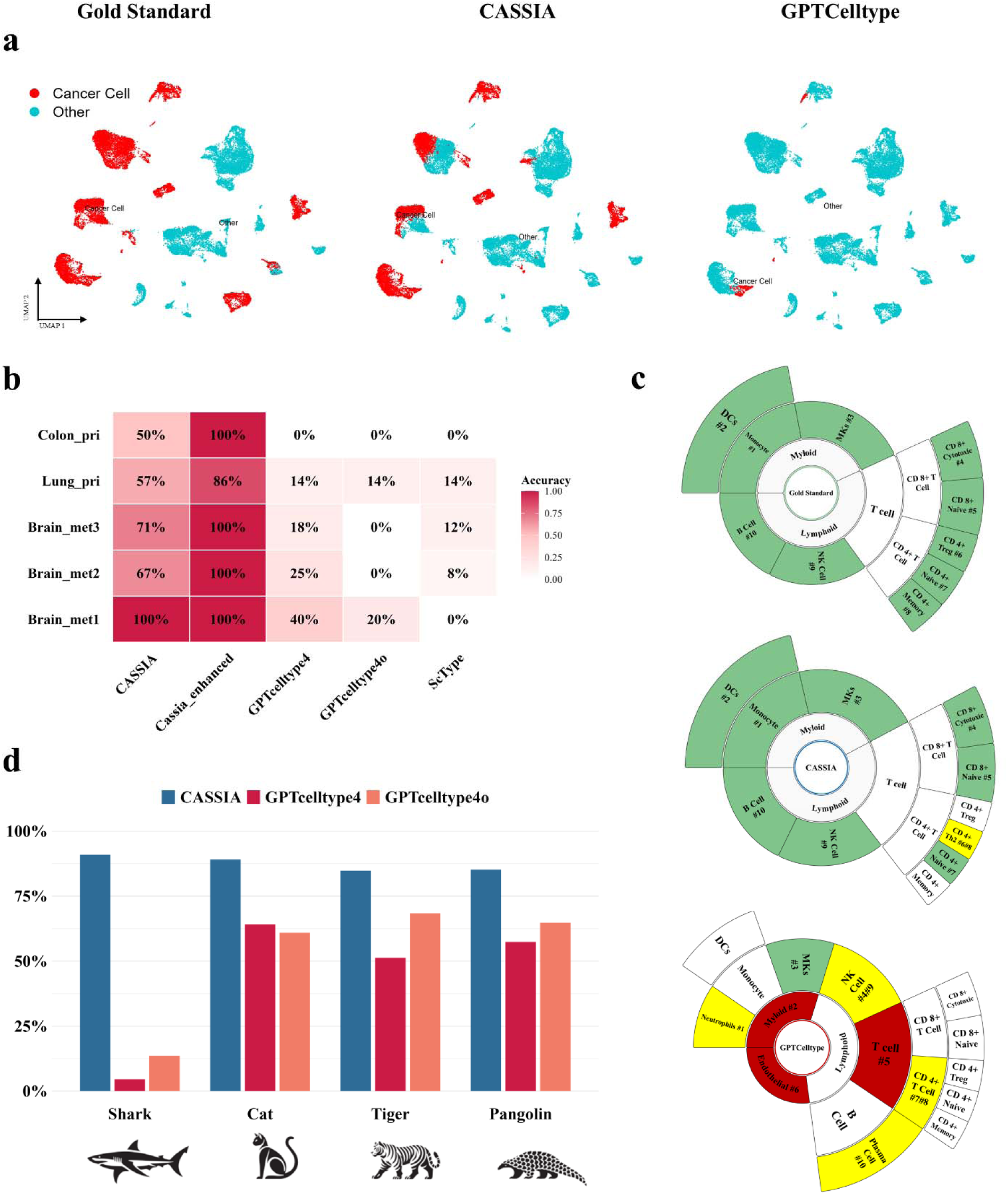
CASSIA outperforms competing methods in annotating complex biological datasets including cancer, immune cells, and rare species. **(a)** UMAP visualizations of cancer cell classifications in brain metastasis samples. Left: Gold standard annotations showing cancer cells (red) and non-cancer cells (blue). CASSIA annotations closely match the gold standard (middle). GPTCelltype4 annotations show significant misclassification of cancer cells as non-cancer (right). **(b)** Heatmap quantifying tumor cell detection accuracy across five cancer datasets (rows) for five annotation methods (columns). CASSIA achieves 57-100% accuracy, while CASSIA-enhanced reaches 86-100% accuracy. Other methods (GPTCelltype4, GPTCelltype4o, ScType) show substantially lower performance (0-40%). **(c)** Radial plots comparing immune cell subtype classification in PBMC samples. Gold standard annotations (top) compared with CASSIA (middle) and GPTCelltype4 (bottom). Color-coding indicates annotation accuracy: green (correct), yellow (partially correct), and red (incorrect). CASSIA accurately identifies most immune cell subtypes and activation states, while GPTCelltype4 shows substantial misclassifications (red sectors). **(d)** Bar chart comparing annotation performance across non-model organisms. CASSIA (blue) consistently achieves >85% accuracy across shark, cat, tiger, and pangolin datasets, while GPTCelltype4 (red) and GPTCelltype4o (orange) show variable and significantly lower performance, particularly in evolutionarily distant species like shark.

We next examined whether this robust performance extends to complex immune cell landscapes, which present distinct annotation challenges due to their hierarchical differentiation patterns and functional plasticity. We considered the PBMC68k dataset which contains major peripheral blood mononuclear cell populations representing a generic immune cell landscape, and the ProjectTILs dataset, a specialized manually curated reference capturing subtle transcriptional differences between T cell functional states. For these immune cell datasets, CASSIA improved average annotation accuracy by 27% over the next best performing method (Figure 2c). While most existing methods perform reasonably well at broad cell type assignments—such as distinguishing T cells from B cells—CASSIA stands out for its ability to preserve high accuracy when annotating fine-grained functional states. Notably, it is the only method that consistently resolves subtle distinctions among naive, memory, and exhausted T cell subsets (Figures 4c and Supplementary Figure 1).

Having demonstrated CASSIA’s capabilities in annotating cancer and specialized immune cell populations, we next sought to determine whether its annotation framework could extend beyond well-characterized human and mouse systems to evolutionarily diverse species with limited reference data. Specifically, we tested CASSIA on datasets from diverse vertebrate species including cartilaginous fish (shark) and several mammals (pangolin, domestic cat, and tiger). CASSIA demonstrates robust cross-species annotation capabilities, correctly identifying 66 of 79 cell subtypes in tiger, 57 of 64 in domestic cat, 20 of 22 in shark, and 41 of 54 in pangolin, increasing annotation accuracy over other methods by 14–77% (Figure 4d). These results further demonstrate CASSIA’s ability to transfer knowledge across phylogenetically diverse species despite the lack of comprehensive reference databases.

### CASSIA’s provides robust annotation-specific quality scores

LLMs almost always provide answers to queries, and they typically do so with uniform confidence. In addition, while generally accurate, LLMs are known to hallucinate, even in recent models like GPT-4o^21,22^. To address these concerns, guidelines for the safe and effective use of LLMs in practice strongly suggest that results be cross checked and validated for accuracy^23,24^. To address this need and thereby allow users to distinguish hallucinations and/or low-quality annotations from high-quality ones, CASSIA employs a two-tiered quality assessment framework. Specifically, CASSIA reports annotation-specific quality scores ranging from 0-100. As the quality score is derived from a single run of CASSIA, an optional Consensus Similarity (CS) score can also be obtained to quantify agreement across multiple CASSIA runs (the CS score is optional as it requires additional computational time due to multiple CASSIA runs; see Methods).

To evaluate CASSIA’s quality assessment framework, we obtained quality scores for 500 cell types across five reference datasets: GTEx, Tabula Sapiens (TS), Human Cell Landscape (HCL), Mouse Cell Atlas (MCA), and Azimuth. For the more computationally intensive CS score calculation, we randomly sampled a total of 100 cell types, drawing 20 cell types from each of the five datasets. As shown in Figure 5a, there is a significant association between quality scores and annotation correctness. Annotations with scores below 75% were predominantly partially or fully incorrect, with a specificity of 0.769 - indicating that low scores reliably flag uncertain or erroneous annotations. The 75% cutoff was selected through comprehensive threshold testing to optimize overall classification performance across precision, recall, and specificity (see Methods). Scores between 75% and 90% reflect intermediate confidence, while scores above 90% are strongly associated with correct classifications. Figure 5b illustrates the relationship between CS scores and annotation accuracy. While the difference in CS scores between partially correct and fully correct annotations was not statistically significant, incorrectly annotated cell types exhibit significantly lower CS scores compared to both of the other categories. This finding suggests that CS scores are particularly effective at identifying incorrect annotations.

**Figure 5:**
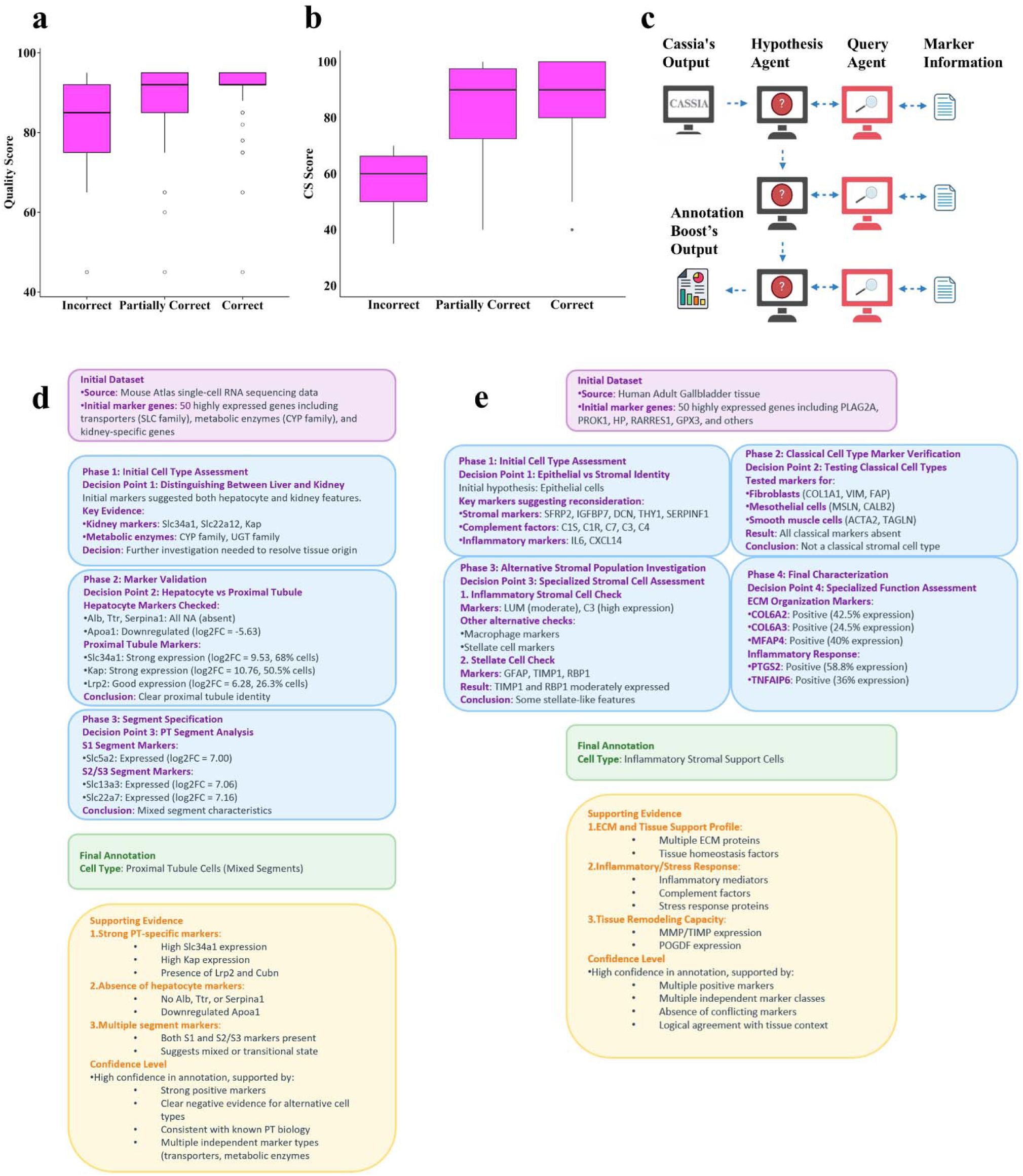
CASSIA’s quality assessment framework provides informative and actionable annotation scoring. **(a)** Box plots demonstrating the relationship between CASSIA’s quality scores and annotation accuracy. Scores are higher for correct annotations, intermediate for partially correct annotations, and lower for incorrect annotations, confirming the diagnostic value of these quality metrics (Kruskal-Wallis test, p<0.01). **(b)** Box plots showing CASSIA’s confidence scores (CS) stratified by annotation accuracy. CS scores effectively discriminate between correct or partially correct (median ∼90) and incorrect annotations (median ∼60), providing reliable confidence estimation. **(c)** Schematic illustration of CASSIA’s Annotation Boost framework, which can be used to refine low scoring annotations. The system works iteratively: the Hypothesis Agent generates initial hypotheses based on CASSIA’s output, then the Query Agent identifies marker genes for validation and retrieves relevant information from FindAllMarkers output files. Results are fed back to the Hypothesis Agent to generate refined hypotheses. This cycle continues through multiple iterations, with the final output from Annotation Boost providing enhanced cell type classifications. Annotation Boost agent reports for challenging cell types including proximal tubule cells from kidney **(d)** and inflammatory stromal support cells **(e)**. Each report presents a structured analytical workflow including: initial dataset overview and highly expressed markers; multi-phase decision points with evidence-based reasoning; marker validation steps with expression thresholds; final annotation with comprehensive supporting evidence; confidence assessment with positive/negative markers; key biological implications of the identified cell type. The reports demonstrate CASSIA’s transparent, systematic approach to complex cell type identification with explicit reasoning chains.

Overall, CASSIA’s annotations matched the GTEx gold-standard annotations on over 95% of the 106 cell types. Quality scores exceeded 85% on 103 of the 106 cell types and surpassed 90% on 86 of the 106 cell types. Other datasets also showed good alignment, though not as strongly, with a specificity of 0.768. As discussed in the next section, this reduction in specificity is likely due at least in part to errors in the gold standard. Taken together, CASSIA’s quality and CS scores provide the user with an informative assessment of annotation-specific confidence, with CS scores being particularly valuable for identifying incorrect annotations.

### The Annotation Boost agent refines low-scoring annotations to improve accuracy

When a cell type is returned with low quality and/or CS scores, the Annotation Boost agent may prove useful in refining the annotation. Unlike the default workflow which relies only on ranked marker lists, the Annotation Boost agent leverages all statistical metrics from the full FindAllMarkers file (including p-values, percentage expression, and log2 fold changes) to generate and test specific hypotheses about cell identity. As illustrated in Figure 5c, the Annotation Boost agent employs an iterative query-and-verify approach, with each cycle refining the annotation hypothesis until convergence or a maximum iteration limit. We considered the 42 annotations (out of 534) having low quality scores (≤75%) across our benchmark datasets; 27 were incorrectly annotated by CASSIA and 15 were correctly annotated. When applied to these 42 annotations, the Annotation Boost agent successfully corrected 24 of the 27 (89%) previously incorrect annotations while preserving all 15 annotations that were originally correct. This improvement was consistent across all evaluated datasets, with the most substantial gains observed in complex tissues with closely related cell types.

A representative example is shown in Figure 5d for a collection of proximal tubule cells. CASSIA, GPTCelltype, and SingleR incorrectly classify these cells as hepatocytes, but CASSIA’s low quality score (75%) suggests further attention. We applied the Annotation Boost agent, which identified a tissue context mismatch and ultimately provided a detailed annotation of “proximal tubule cells (mixed segments)”. The sequential reasoning pattern of the agent, visible in the chat history excerpts in Figure 5d and 5e, demonstrates how CASSIA systematically builds evidence for its revised annotation by identifying and then querying the expression patterns of additional marker genes that were not in the initial ranked list.

Another notable refinement involved cells labeled as “stromal cell_PLA2G2A high” that were misidentified as epithelial cells by CASSIA and all other methods. Given CASSIA’s quality score for this cell type was 75%, the Annotation Boost agent was applied and correctly identified these cells as “inflammatory stromal support cells” by first ruling out epithelial characteristics through additional stromal marker identification, then conducting a specialized functional assessment that revealed the inflammatory state associated with elevated PLA2G2A expression (Figure 5e). These examples demonstrate how the Annotation Boost agent extends CASSIA’s capabilities beyond initial annotation and facilitates the refinement of challenging annotations through expert-like reasoning processes.

### CASSIA’s quality assessment framework identifies heterogeneous cell populations and errors in gold standard datasets

To further evaluate CASSIA’s quality assessment framework, we evaluated cell types with low quality scores as well as those with high quality scores that contradicted the gold standard annotations. First, we considered two datasets - the small intestine and eye TS datasets- as these had low quality scores on average. For these datasets, cell type identification is particularly challenging due to poorly separated clusters and unbalanced cell counts. In the small intestine dataset, we identified annotations with low CS scores, including goblet cells (75%) and mature enterocytes (30%). Further investigation revealed the goblet cell cluster contained mixed cell types while the mature enterocyte cluster showed an unusually high mitochondrial ratio (>60%) (Figure 6a). Similar patterns were observed in the TS eye dataset (Supplementary Note 2). These analyses demonstrate that low CS scores are often derived from low-quality cell populations or heterogeneous populations that require additional subclustering.

**Figure 6:**
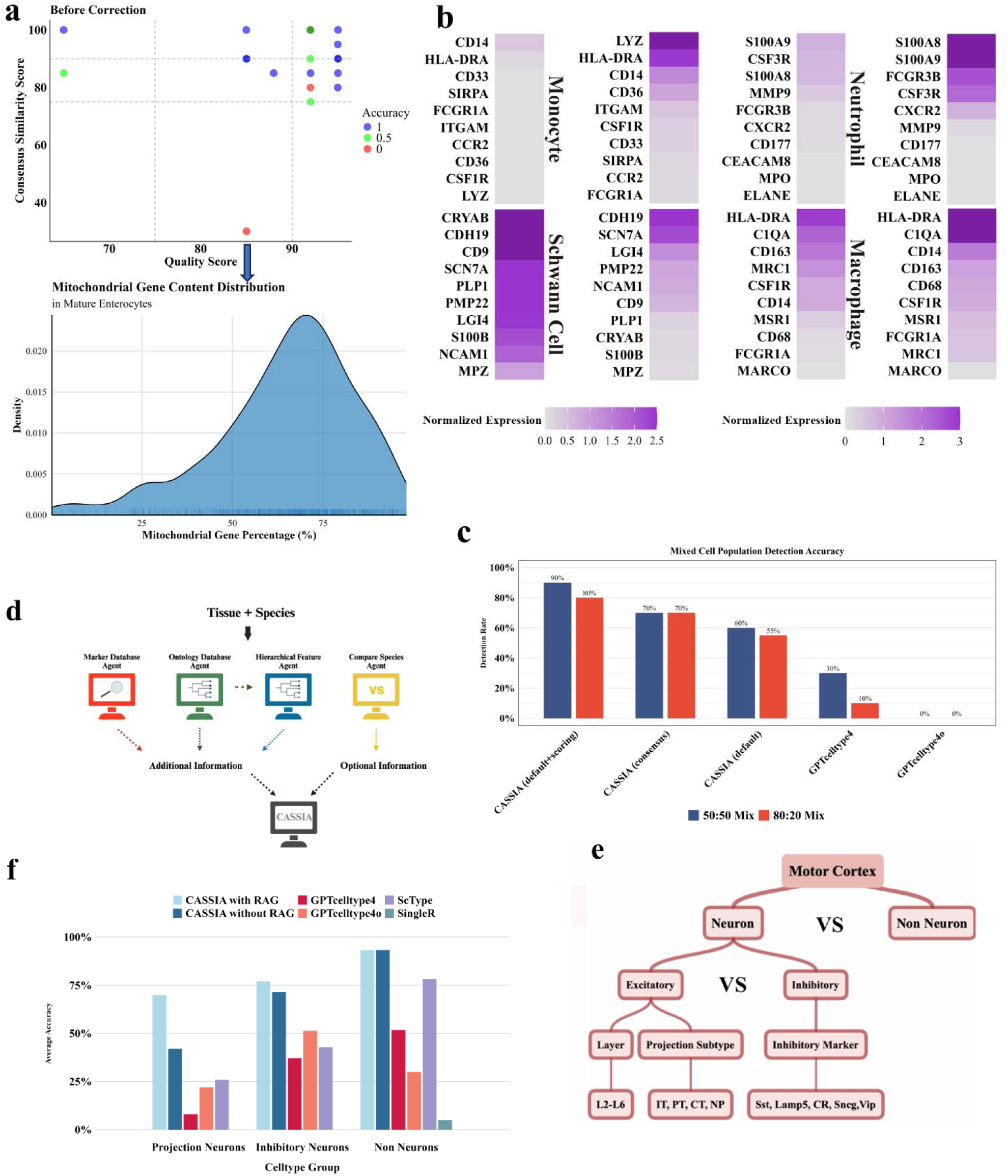
CASSIA’s quality assessment framework identifies gold standard annotation errors while the RAG agent enhances performance for challenging cell types. **(a)** Scatter plot of quality scores and CS scores for cell type annotations in the TS small intestine dataset, colored by annotation accuracy; the mature enterocyte cluster has an abnormally low CS score near 30% (top). Distribution of mitochondrial gene content in the mature enterocytes cluster. Most cells show elevated mitochondrial gene ratios (>60%), indicating poor quality data which is reflected in the low CS score (bottom). **(b)** Heatmaps of marker expression comparing CASSIA’s annotations with gold-standard annotations. The leftmost column considers cell types that CASSIA identifies as Schwann cells, but the gold standard identifies as monocytes. Shown are marker gene expression averages in those cells for canonical Schwann cell markers and monocyte markers. The high expression of Schwann cell markers and minimal expression of monocyte markers strongly suggests that the gold-standard annotation of monocytes is incorrect. The second column is a positive control, showing expression of those same markers in cell types that both CASSIA and the gold standard agree are monocytes (top) or Schwann cells (bottom); we see relatively high expression for at least some of these markers, showing that the markers together do in fact represent the cell type. The third column shows cells that CASSIA identifies as macrophages, but the gold standard identifies as neutrophils. Shown are marker gene expression averages in those cells for definitive macrophage markers and neutrophil markers. The high expression of macrophage markers and minimal expression of neutrophil markers suggests that the gold-standard annotation is incorrect. The last column is a positive control, showing expression of those same markers in cell types that both CASSIA and the gold standard agree are neutrophils (top) or macrophages (bottom). These examples demonstrate CASSIA’s ability to detect annotation inconsistencies. **(c)** Quantification of mixed cell type detection capabilities in synthetic mixtures. Bar chart comparing detection accuracy across different methods for both 50:50 mixtures (blue bars) and 80:20 mixtures (red bars). CASSIA default relies solely on CASSIA’s report of mixed cell types and examination of sub-cell types. CASSIA consensus (running default CASSIA 5 times and applying majority voting) considers mixed cell types detected and applies CS score thresholds. Both CASSIA methods substantially outperform GPTCelltype approaches across both mixture scenarios. **(d)** Schematic of CASSIA’s Retrieval-Augmented Generation (RAG) pipeline. The system integrates specialized agents accessing the CellMarker2.0 database for validated markers and the Owlready ontological database for tissue-specific hierarchies. The Hierarchical Feature agent identifies hierarchical patterns that effectively categorize cell types into functionally related groups; and the cross-species comparison agent analyzes evolutionary conservation between the species of interest and reference species to provide guidance on marker interpretation accounting for species-specific differences. These components provide contextual knowledge to enhance CASSIA’s default annotation pipeline. **(e)** Hierarchical classification schema for mouse motor cortex neurons, illustrating the multi-level annotation challenge from basic neuron identity to excitatory/inhibitory classification, layer specification, projection subtype determination, and inhibitory marker profiling. **(f)** Comparative performance in annotating neuronal subtypes in mouse motor cortex. CASSIA with RAG (light blue) demonstrates superior accuracy across projection neurons (∼70%), inhibitory neurons (∼75%), and non-neurons (∼95%) compared to CASSIA without RAG and other methods demonstrating that the RAG enhancement improves identification of specialized neuronal subtypes.

We also investigated high-confidence annotations that contradicted established reference labels. Specifically, we first examined 15 annotations (out of over 500 total) where CASSIA’s quality scores exceeded 90%, but CASSIA’s annotation did not match the gold standard. For each annotation, we built an evaluation system using three large language models (LLMs). Each LLM was given the marker gene profile of the cell type and asked to determine which label—CASSIA’s prediction or the gold standard—the profile more closely resembled, assigning a score to each option. When all three LLMs consistently favored CASSIA’s annotation, we manually analyzed and visualized the key classical markers.

Of the 15 annotations reviewed, 11 (73.3%) agreed with CASSIA’s annotation, while only 1 (6.7%) disagreed. The remaining 3 cases (20%) yielded conflicting or inconclusive results across the different LLMs (see Supplementary Note 2c). Figure 6b shows two representative examples where the three LLMs agree with CASSIA’s annotations: cells identified by CASSIA as “Schwann cells” but labeled in the reference as “monocytes,” and cells identified as “macrophages” but labeled in the reference as “neutrophils.” To investigate these discrepancies, we identified canonical cell type markers with high discriminatory power for each contested cell type pair (monocytes vs. Schwann cells; neutrophils vs. macrophages). Figure 6b shows that the marker gene expression profiles strongly support CASSIA’s annotations. Additional examples are provided in Supplementary Figure 4 and Supplementary Data 9.

### CASSIA identifies mixed cell types

To evaluate CASSIA’s ability to recognize mixed cell types, we synthetically generated input profiles by combining the top 50 marker genes from two distinct cell types in defined ratios (50:50 and 80:20). We compared results from running CASSIA in both default mode and consensus mode with 5 repetitions, the latter providing CS scores, consensus cell type annotations, and potential mixed cell type identifications.

In the balanced 50:50 mixing scenario, default CASSIA directly identified mixed populations in 6 out of 10 cases. An additional 3 cases were flagged by CASSIA’s scoring agent as potentially mixed, yielding an overall detection accuracy of 90%. The consensus mode performed similarly, detecting one additional mixed situation (7/10), though it misidentified the specific mixed cell types in this additional case. In contrast, GPTCelltype4 failed to detect any mixed populations, and GPTCelltype4o identified only 3 out of 10 mixed cases (Figure 6c).

In the more challenging 80:20 mixing scenario, default CASSIA directly identified mixed populations in 11 out of 20 cases, with the scoring agent detecting 5 additional mixed profiles. This resulted in an 80% overall detection accuracy. The consensus mode demonstrated superior performance, correctly identifying mixed populations in 14 out of 20 cases. Both default CASSIA and consensus mode CASSIA achieved an average dominant cell type annotation accuracy of 0.9, correctly identifying the cell type contributing 80% of the markers. By contrast, GPTCelltype4 and GPTCelltype4o showed substantially lower performance, detecting only 0 and 2 mixed populations, respectively, and achieving average dominant cell type annotation accuracies of 0.35 and 0.45.

### CASSIA’s RAG agent enhances annotation of complex tissues

While CASSIA performs well on standard datasets with its core functionality, some tissues with complex hierarchical organization require additional domain-specific knowledge for optimal annotation. Toward this end, CASSIA employs a Retrieval-Augmented Generation (RAG) agent that leverages tissue-specific markers from external databases and ontologies to refine annotations and accurately subdivide major cell types into specialized ones.

We here provide an example using the RAG multi-agent pipeline to enhance annotation of the mouse motor cortex in the Azimuth dataset (an overview of the pipeline is given in Figure 6d). Neurons make up the largest cell type in this dataset, and the general class of neurons is relatively straightforward to annotate. However, a detailed classification requires annotation of major type (excitatory, inhibitory), as well as numerous subtypes (local vs. projection, and within projection intratelencephalic (IT), extratelencephalic (ET), corticothalamic (CT), or near projection (NP)), inhibitory subtype (Pvalb+, SST+, VIP+, and Lamp5+), and layer location (2-6). Figure 6e illustrates this hierarchical organization of neuronal subtypes in the motor cortex. We use the RAG agent here to provide additional background information to CASSIA related to cell types and their corresponding markers. In this analysis, the Marker Database agent identified 43 markers based on seven related cell types including glutamatergic neurons, GABAergic neurons, astrocytes, dopaminergic neurons, oligodendrocytes, microglia, and vascular cells. The Ontology Database agent identified and filtered ontology trees to ensure specificity to the mouse cortex; the initial trees identified were rooted by neurons, glial cells, vascular cells, and meningeal cells and then restricted to only contain mouse cortex cells, resulting in four trees overall. The Hierarchical Feature agent performed a hierarchical analysis on the four ontology trees selected from the second module to identify key discriminative features and associated markers. For example, in the neuron-rooted tree, there are three sublevels that discriminate features within the cortex ontology: inhibitory vs. excitatory, laminar location, and projection patterns. Markers associated within each of the sublevels are then identified by the RAG agent and provided to CASSIA. As shown in Figure 6f, CASSIA with RAG is the only method capable of precise annotation of neuronal subtypes.

## Discussion

Single-cell RNA sequencing has revolutionized our understanding of cellular heterogeneity, yet accurate cell type annotation remains a critical bottleneck. CASSIA addresses this challenge through a multi-agent LLM framework that provides automated, accurate, and interpretable cell annotations. The benchmarking studies presented here demonstrate that CASSIA improves annotation accuracy in large benchmark datasets, and also in more complex datasets from studies of cancer, immunology, and rare species. In addition, CASSIA’s quality assessment framework provides annotation-specific quality scores that identify low-quality annotations which can be investigated manually, or via the Annotation Boost agent.

Despite these advances, limitations remain. CASSIA’s performance depends on marker gene quality, which may be challenging to identify in datasets with poorly defined clusters or continuous trajectories; and the initial marker calculation process is computationally intensive and time-consuming, particularly for large datasets. Future directions include deeper integration with Seurat and Scanpy to streamline the workflow from raw data to marker identification to annotation. A key advancement would be developing adaptive marker selection algorithms that dynamically determine the optimal number of markers to use based on cluster characteristics, rather than applying a fixed threshold of 50 markers. This approach would include new algorithms to rapidly identify the most representative markers for each cluster, potentially reducing computational burden while maintaining or improving annotation accuracy. Additional directions include extending capabilities to iterative clustering and trajectory analysis, and incorporating complementary results (such as from gene set enrichment analysis) to further enhance annotation precision.

In conclusion, CASSIA leverages the reasoning capabilities of LLMs for robust validation, interpretability, and quality assessment to provide accurate and transparent cell type annotation across diverse biological contexts. Importantly, CASSIA provides a user with the logical reasoning behind each annotation along with annotation-specific quality scores, reducing the black-box^25^ nature of LLM-based approaches and guarding against hallucinations. Ultimately, CASSIA’s prompt-based nature lowers the computational expertise needed to conduct single-cell annotation, while also providing more biological-specific expertise through the RAG agent. Taken together, CASSIA increases both the accuracy and accessibility of cell type annotation in practice.

## Data Availability

The dataset(s) used here are available as follows: Raw RNA sequencing data generated in this study have been deposited in the Gene Expression Omnibus (GEO) repository under accession number (pending). All other datasets used are publicly available and are listed in Supplementary Table 1.

## Software Availability

- Project name: CASSIA
- Project home page: https://github.com/ElliotXie/CASSIA^22^
- Archived version: CASSIA 1.0.0
- perating system(s): Platform-independent
- Programming language: Python, R
- License: MIT License

## Author contributions

EX conceived the study and led the method development, implementation, validation, and case study analyses. CK co-led the project. LC contributed to the method development and developed the RAG agent. LC, YC, and JL contributed to case study analyses. MD collected the brain tumor samples and assisted with interpretation of results; JS processed the brain tumor samples for scRNA-seq profiling, contributed to method development, and assisted with interpretation of results from all case study analyses. CK, EX, LC, and JS wrote the manuscript.

## Acknowledgments

This work was supported by NIH GM102756 (CK), NIH K08NS092895 (MD), and the IU Value Research grant (MD).

## Methods

CASSIA is a multi-agent LLM consisting of an onboarding platform and five interconnected LLMs: an annotation agent, a validation agent, a formatting agent, a quality scoring agent, and a reporter agent. Optional LLMs are also available for subclustering, uncertainty quantification, and annotation boosting. For applications requiring highly detailed annotations, CASSIA includes a RAG agent. Each agent is described in detail below.

### CASSIA implementation

We provide CASSIA as an R package, a Python package, and a web-based user interface (https://www.cassiacell.com/ ^23^). Both programming packages support all CASSIA functionalities including optional agents, with the default pipeline executable via a single command. The web interface implements the default pipeline only.

### Onboarding

A user provides information to an Onboarding platform by specifying species, tissue type, and a collection of markers associated with cell subtypes within that tissue, if known. Most analyses will use markers from *FindAllMarkers* in Seurat or *tl.rank_genes_groups* in Scanpy, although markers can be manually added if desired. We suggest 50 markers be used if available (see the Identification of Marker Genes section below). Any information associated with experimental conditions, interventions, or other sample-specific information may be provided, and will be used to create the user prompt given to the Annotator agent. The specific user prompt reads: “Your task is to annotate a single-cell [species] dataset from [tissue type]. Please identify the cell type based on this ranked marker list: [marker list]. Below is some additional information about the dataset: [additional information].” In some cases (e.g. multiple mixed cell types from an atlas), tissue type is not known. A tissue blind version can also be used (see Supplementary Note 3).

### Annotation

The Annotator agent performs a comprehensive annotation of the single-cell data. To minimize hallucinations and prevent oversimplification of complex biological concepts, we implemented multiple prompt engineering techniques: a zero-shot chain-of-thought approach that structures the model’s reasoning to follow the same systematic analytical steps a professional computational biologist would take^26,27^, role specification where the Annotator agent acts as a ‘professional computational biologist with expertise in single-cell RNA sequencing,’ and audience framing as ‘an expert in the field.’ These role-playing strategies have been shown to improve the likelihood of receiving detailed and technically accurate responses^28^. Finally, we included a hypothetical reward statement: ‘you will be rewarded $10,000 if you do a good job’ as such reward modeling in LLMs has been shown to prime the model for maximum effort and attention to detail^29^.

Taken together, the system prompt to the Annotator Agent reads: “You are a professional computational biologist with expertise in single-cell RNA sequencing (scRNA-seq). A list of highly expressed markers ranked by expression intensity from high to low from a cluster of cells will be provided, and your task is to identify the cell type. You must think step-by-step, providing a comprehensive and specific analysis. The audience is an expert in the field, and you will be rewarded $10000 if you do a good job. Steps to follow:

1. List the Key Functional Markers: Extract and group the key marker genes associated with function or pathway, explaining their roles.
2. List the Key Cell Type Markers: Extract and group the key marker genes associated with various cell types, explaining their roles.
3. Cross reference Known Databases: Use available scRNA-seq databases and relevant literature to cross reference these markers.
4. Determine the Most Probable General Cell Type: Based on the expression of these markers, infer the most likely general cell type of the cluster.
5. Identify the Top 3 Most Probable Sub Cell Types: Based on the expression of these markers, infer the top three most probable sub cell types. Rank them from most likely to least likely. Finally, specify the most likely subtype based on the markers.
6. Provide a Concise Summary of Your Analysis.

Always include your step-by-step detailed reasoning. You can say “FINAL ANNOTATION COMPLETED” when you have completed your analysis. If you receive feedback from the validation process, incorporate it into your analysis and provide an updated annotation.”

Once FINAL ANNOTATION COMPLETED is reached, annotations are provided to the Validator agent to ensure accuracy and consistency.

### Validation

The Validator agent is designed to iteratively cross check the results provided by the Annotator agent, forming a feedback loop that significantly enhances the robustness of the final annotations. This iterative design is inspired by self-verification techniques, which allow for critical evaluation and continual refinement of annotations^30^. The Validator agent primarily ensures marker and cell type consistency, verifying that the key markers identified by CASSIA are present in the provided marker list and accurately represent the identified cell type. This step is crucial to confirm that the markers used for annotation are both appropriate and reliable. If the validation fails, the Validator agent provides detailed feedback and sends the results back to the Annotator agent for revision. This process iterates up to three times to ensure the highest quality annotation. Regardless of whether validation finally passes or fails, the results are then forwarded to the Formatting agent for further processing. The full system prompt for the Validator agent is provided in Supplementary Note 3.

### Formatting

A formatting agent was implemented to process annotation outputs from the previously mentioned two agents. The agent operates in two distinct states: a standard mode that extracts and structures validated cell type annotations including mixed population information, and a diagnostic mode that activates upon validation failure to analyze error sources and generate recommendations for resolution. The full system prompt is provided in Supplementary Note 3.

### Quality Scoring

The Scoring agent functions as an objective evaluator that analyzes the complete annotation conversation history to generate a quantitative assessment score (0-100%). It focuses on scientific accuracy of the annotations along with balanced use of multiple markers where high-ranked markers contribute most to the annotation and no single marker drives the annotation. The full system prompt for the Quality Scoring agent is provided in Supplementary Note 3.

To establish thresholds for the quality scores, we implemented a weighted cost optimization approach defined as Total cost = cw x FPJ + FN where FP represents false positives (correct annotations incorrectly flagged as low quality), FN represents false negatives (incorrect annotations misclassified as high quality), and w is the incorrect weight parameter (default = 2). We assigned greater weight to false positives to minimize the risk of unnecessarily flagging correct annotations for manual review. Supplementary Figure 5 shows that the lower threshold minimizing total cost is ∼75% when evaluated on more than 500 cell type annotations from the five benchmark datasets. Given this, annotations with quality scores below 75% are flagged as low-quality. In these cases, the (optional) Annotation Boost agent could be used to improve the annotation.

### Reporter agent

The Reporter agent generates a comprehensive HTML report documenting the complete annotation process, including agent conversations, quality evaluation reasoning, and validation decisions with supporting evidence. The full system prompt is provided in Supplementary Note 3.

### Subclustering Agent

CASSIA provides an optional subclustering agent designed to optimize the analysis of specialized cell populations (e.g., CD4+ T cells cluster). Rather than processing individual subclusters separately, this agent analyzes all subclusters within a specific cell population simultaneously. This concurrent analysis strategy improves computational efficiency, enabling better detection of subtle phenotypic differences between closely related cell states. The full system prompt is provided in Supplementary Note 3.

### Uncertainty quantification

CASSIA provides an optional feature to assess annotation reliability by quantifying consensus across multiple (defaults to 5) runs. The Consensus Similarity (CS) score described below builds upon recent advances in LLM ensemble methods, which suggest that multiple iterations of a model or cooperation between several different models can achieve superior performance through consensus mechanisms^31,32^.

This consensus approach is grounded in probability theory, where running a model multiple times with majority voting significantly improves accuracy. For a model with base accuracy p (e.g., 0.7 or 70%), the probability of getting a correct result after n runs with majority voting follows a binomial distribution.

Specifically, if we need at least k correct answers out of n runs (where k = ceiling(n/2) for majority), the probability is:

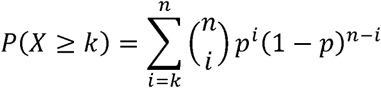

where 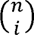 is the binomial coefficient. This mathematical foundation explains why even 5 runs with majority voting can substantially improve reliability. For example, a model with 70% base accuracy achieves approximately 84% accuracy after 5 runs, and nearly 95% accuracy after 15 runs. For models with higher base accuracy (e.g., 80%), the improvement is even more dramatic, approaching 99.9% accuracy after just 20 runs (Supplementary Figure 6).

To ensure consistent comparisons across CASSIA runs, cell type nomenclature is standardized using two complementary approaches. The first approach applies basic text normalization (converting to lowercase, replacing hyphens with spaces, removing trailing ‘s’), then maps each normalized label to standardized Cell Ontology terms via the CL API¹². This mapping matches cell types to their formal ontological definitions, ensuring consistent terminology. After unifying cell type names, we employ a consensus similarity (CS) score to measure the similarity across the *A* CASSIA runs. Recall that the Annotator agent identifies the most probable general cell type and the three most probable sub-cell types, ranked from most likely to least likely. Letting *R* denote the set of all annotation results, each result *r ε R* is a pair (*g, s*) where *g* represents the general cell type and *s* represents the most likely sub-cell type. Then

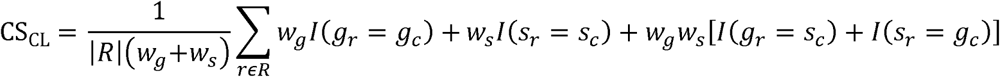

where *g*_C_ is the most frequent general cell type in *R*; s_C_ is the most frequent cell sub-type in *R*; *w_g_* and *w*_S_ specify the weights given to matches of the general cell type and cell sub-type, respectively. *I(x)* denotes the indicator function where *I(x)* =1 if *x* is true and 0 otherwise; |*R*| denotes the number of annotations in *R*. We use CS_CL_ to emphasize the fact that this CS score was calculated using terms unified by the Cell Ontology CL API. The CS_CL_ score takes into account both the general and sub-cell types, giving more weight to matches in the general type while still considering sub-type matches and potential type inversions.

To address potential API mapping inconsistencies, a second approach employs an LLM-based unification agent that harmonizes synonyms and slight variations in cell type nomenclature. A CS_LLM_ based score is calculated based on this unification. Finally, to enhance robustness and mitigate potential errors from unification steps, a consensus agent independently estimates the consensus general cell type and subtype. Taken together, three measures of consensus are obtained (CS_CL_, CS_LLM_, and the consensus agent’s estimated consensus score). The minimum score is taken as the final CS score. Annotations with a CS score below 75% are flagged as uncertain and recommended for further validation using the Annotation Boost agent.

### Annotation Boost agent

CASSIA provides an optional annotation agent that can be used to improve low-scoring annotations (below 75%). Unlike the default workflow which uses only ranked marker gene lists, the Annotation Boost agent leverages comprehensive statistical metrics from the *FindAllMarkers* file uploaded to the Onboarding platform (e.g. p-values, percentage expression in groups, and log2 fold changes) and examines expression patterns across all genes. Based on results from the Annotation agent, it generates testable hypotheses and systematically queries gene expression data to assess each hypothesis. This process continues iteratively until either a high-quality annotation is achieved, or the maximum iteration limit (defaults to 5) is reached. Further details and the full system prompt are provided in Supplementary Note 3.

### Retrieval-Augmented Generation (RAG) pipeline

For cell types with markers that are mentioned relatively infrequently in the literature, and therefore not well represented in the LLM that CASSIA’s annotator agent uses (defaults to GPT4o), an optional multi-agent RAG pipeline can be useful. The RAG pipeline consists of multiple sequential agents that leverage established cell marker databases and biological ontologies to generate comprehensive marker profiles for tissue-specific cell type identification. Specifically, the Marker Database agent queries the CellMarker database^19^ to identify validated, canonical cell type markers. The agent will generate a conservative list of markers specific to the tissue of interest. Given the markers, which in many cases contain markers with relatively low sensitivity and specificity, the agent further analyzes the marker list to identify markers of broad cell types as well as those representing refined tissue-specific subtypes. The second agent, the Ontology Database agent, leverages the Owlready ontological database^33^ to identify those ontology trees associated with the tissue of interest. Trees are then filtered by the agent to retain only those containing relevant tissue-relevant cell types, eliminating extraneous branches that are not typically present in the tissue of interest. The Hierarchical Feature agent then performs a PCA-like analysis on the ontology trees selected from the second agent to identify key discriminative features. This process identifies hierarchical patterns that effectively categorize cell types into functionally related groups that ultimately inform the cell type annotation. For each identified feature axis, we compile specific markers that can effectively discriminate between cell populations along that axis. For studies involving non-model organisms not directly represented in reference databases, we implemented an optional cross-species comparison agent. This agent analyzes evolutionary conservation between the species of interest and reference species to evaluate marker gene conservation, identify potential inter-species variations, and provide guidance on marker interpretation accounting for species-specific differences. The final component consolidates all generated outputs in the previous four agents into a text format, which is then integrated as additional information into the default CASSIA pipeline. Details on the evaluation procedure, including annotation accuracy assessment and the scoring criteria used in the RAG pipeline testing, are provided in Supplementary Data 2, while further methodological information, including the full system prompt, is available in Supplementary Data 3.

### Fully correct vs. partially correct annotations

We established a hierarchical evaluation framework based on the Cell Ontology tree structure to assess annotation accuracy, classifying annotations as fully correct, partially correct, or incorrect based on their taxonomic distance from reference annotations. An annotation one step away from the reference on the ontology tree is considered partially correct (e.g., predicting “T-cell” for a CD8+ T-cell), while annotations multiple steps removed are classified as incorrect (e.g., predicting “immune cell” for a CD8+ T-cell). For rare or highly specific cell types, we adapted this framework to consider annotations as partially correct when they accurately identify the general cell type category, such as classifying “epithelial cell” as partially correct for Bestrophin 4 (BEST4) + cells. See Supplementary Data 2 for more evaluation examples. A similar, but more detailed, scoring system was used for the RAG agent applied to the mouse motor cortex data (Supplementary Data 2).

### Average accuracy

To calculate average accuracy, cell type annotations are evaluated on a three-tier scale: fully correct annotations are assigned a score of 1, partially correct a score of 0.5, and incorrect a score of zero. Average accuracy for a dataset is the average score across all annotated cell types for that dataset.

### Identification of marker genes

We evaluated CASSIA’s performance across varying marker set sizes (*n* = 10, 30, 50, 70, 100) using the GTEx data, large and small intestine tissues from the Tabula Sapiens (TS), and non-model mammalian samples including domestic cat, tiger, and pangolin. For most datasets, average annotation accuracy plateaued at 50 marker genes (Figure 2d), with minimal improvements observed with larger sets. Based on this analysis, we recommend using 50 marker genes as the optimal balance between computational efficiency and classification accuracy.

### Parallel computing and computational time

CASSIA utilizes Python’s *concurrent.futures* module with *ThreadPoolExecutor* for efficient parallel processing. Cell type analyses are distributed across a user-defined thread pool (default: 10 workers), enabling concurrent annotation of multiple cell types. This parallel architecture scales efficiently, processing each cell type in about 30 seconds per CPU core with pre-prepared marker files. On a standard 8-core computer, CASSIA completes the annotation workflow for typical single-cell datasets with 20 clusters in under 2 minutes.

### Memory management

CASSIA implements selective memory access across its agent network to optimize annotation processing. The annotation and validation agents maintain access to their shared conversation history for iterative refinement, while the scoring agent evaluates the complete dialogue record. For successful annotations, the formatting agent processes only the final validated exchange, whereas for failed cases, it analyzes the full conversation history to generate comprehensive error reports. This targeted memory management strategy ensures efficient information flow while maintaining contextual isolation between distinct cell type analyses.

### Model selection

To evaluate CASSIA’s multi-agent framework across different large language models, we conducted comprehensive benchmarking using three distinct models: Claude 3.5 Sonnet (2024-10-22), GPT-4o (2024-08-06), and LLaMA3.2 90B. We created a benchmark dataset consisting of 100 cell types drawn from five randomly selected tissues within our benchmark datasets to ensure diverse representation. Performance was evaluated against GPTCelltype implementations using both GPT-4 and GPT-4o. Our analysis revealed that CASSIA’s architecture enhances annotation accuracy regardless of the underlying language model. Notably, even when using LLaMA3.2 90B, an open-source model that typically demonstrates lower performance metrics compared to proprietary models, CASSIA achieved superior results compared to GPTCelltype implementations. While Claude 3.5 Sonnet achieved the highest accuracy, we selected GPT-4o as the default model due to its optimal balance between performance and computational efficiency.

### Model implementation and costs

CASSIA supports flexible API integration through three options: OpenRouter API, OpenAI API, or Anthropic API. While direct API connections to OpenAI and Anthropic are supported, the OpenRouter API option enables users to access most of the commercially available large language models without code modifications, including but not limited to GPT-4, Claude, and LLaMA variants. Based on our benchmarking results and cost analysis as of October 2024, we established a tiered recommendation system. GPT-4o serves as the default model, costing approximately $0.02 per annotation ($2.50/M input tokens, $10.00/M output tokens). For applications requiring maximum accuracy, Claude 3.5 Sonnet provides enhanced performance at $0.03 per annotation. For large-scale analyses (>10000 clusters) or cost-sensitive applications, LLaMA3.2 90B offers a cost-effective alternative at $0.003 per annotation through OpenRouter.

### Data preprocessing

Dataset-specific markers were used in most cases, if provided. Exceptions included the shark dataset, where we excluded unmapped genes. Details are provided in Supplementary Table 1. For datasets lacking predefined marker genes, we implemented a standardized analysis pipeline. Raw count matrices were normalized using the *NormalizedData* function in Seurat v5, and differentially expressed genes were identified using Seurat’s *FindAllMarkers* function with parameters set to a minimum cell expression threshold (min.pct) of 0.1, adjusted p-value cutoff of 0.05, and log2 fold change threshold of 0.25.

### Comparative evaluation of existing cell type annotation methods

For comprehensive benchmarking, we compared CASSIA with established annotation methods including GPTCelltype (version 1.0, using GPT4 and GPT4o), ScType^11^ (version 1.0), SingleR^6^ (version 2.4.1), CellTypist^34^ (version 1.6.3), and scCATCH^10^ (version 3.2.2). Full analysis results are detailed in Supplementary Data 4. The optional agents in CASSIA were not used for benchmarking and so all CASSIA benchmarking results demonstrate performance of CASSIA’s default model. As described in the text, the uncertainty quantification agent was used for further analysis of the TS intestinal dataset; the RAG agent was used to analyze the neuronal dataset; and the subclustering agent was applied to the ProjectTILs dataset.

Analysis with each package followed the author’s standard workflow. A few methods required us to specify preferences as follows: for analysis with GPTCelltype, we set the marker size parameter to 10, as recommended; tissue specificity was configured according to each dataset’s context. Analysis with ScType was restricted to tissues with available marker references; gold-standard annotations were used as reference cluster labels. We used majority voting for cluster annotations within SingleR. Special considerations were required for two key datasets: the Human Cell Landscape (HCL) analysis was restricted to marker-based approaches due to annotation-metadata discrepancies, while the mixed-tissue nature of the Mouse Cell Atlas (MCA) limited our comparison to CASSIA and GPTCelltype.

### Immune system analyses

We evaluated CASSIA using two immune datasets. For the PBMC68k dataset, we applied the default CASSIA workflow. For the ProjectTILs dataset, we created a challenging test case for the subclustering agent by integrating specialized human CD4+ and CD8+ T cell reference datasets that capture T cells in various functional states. Since the dataset contains exclusively T cells with subtle state-specific transcriptional differences rather than distinct cell types, it provided an ideal scenario to evaluate CASSIA’s subclustering agent for fine-grained annotation, aligning with our hierarchical annotation design.

### Cancer dataset analyses

We evaluated CASSIA’s performance on cancer datasets, focusing on three aspects: annotation accuracy in cancer-altered transcriptional states, cancer cell detection, and metastatic signal identification. CASSIA successfully identified cancer-specific clusters across all datasets. In three metastasis datasets, CASSIA detected and correctly classified carcinoma cells in non-epithelial anatomical sites (brain and lymphoid tissues). This tissue-discordant detection of carcinoma cells, which originate exclusively from epithelial tissues, demonstrates CASSIA’s ability to identify metastatic signatures through the recognition of cells in developmentally distinct anatomical locations.

### Mixed population simulation analysis

To systematically evaluate CASSIA’s performance in detecting and annotating mixed cell populations, we developed a controlled benchmarking framework using the GTEx lung dataset, selected for its high-quality cell type annotations and well-defined clusters. We generated synthetic mixed populations through a structured marker-mixing strategy. For each simulation, we selected pairs of distinct cell types and created hybrid marker profiles using two mixing ratios: balanced (50:50) and dominant (80:20). In the balanced scenario, we randomly selected 50% of markers from each cell type and interleaved them in alternating order (AℒBℒAℒBℒ…AnℒBnℒ). For the dominant scenario, we selected 80% of markers from one cell type and 20% from the other, maintaining the same interleaving pattern. We evaluated these synthetic populations using both CASSIA, GPTCelltype4, and GPTCelltype4o. In the balanced mixing condition, we assessed each method’s ability to detect the presence of mixed populations. For the dominant scenario, we evaluated both the detection of mixed populations and the accurate identification of the predominant cell type (defined by 80% marker representation). This framework provides a rigorous assessment of each method’s capacity to handle complex cellular compositions while maintaining annotation accuracy.

### Brain metastases tissue collection and scRNA-seq profiling

Tissue samples from patients with confirmed brain metastatic lesions were obtained during the published clinical trial duration of NCT03398694 and obtained and processed as detailed in Shireman *et al*. 2024^35^. Briefly, patients with up to 4 brain metastatic lesions with at least one symptomatic lesion were consented and enrolled in the clinical trial which was approved and monitored by the IU Simon Cancer Center institutional review board and data safety monitoring committee. Confirmation of brain metastatic lesion was done based on MRI imaging, IHC analysis under pathologist supervision, and previous known diagnosis of systemic primary cancer. For this study, patients with primary diagnosis of non-small cell lung cancer and an associated brain metastatic lesion needing resection were taken to the OR and a tissue specimen was obtained by the consulting surgeon. The resulting tissue was dissociated and subject to single-cell sequencing analysis using the 10× chromium method according to manufacturer’s protocol. For single-cell annotation of tumor tissue, gold standard reference for presence of tumor cells and categorization of brain metastasis or primary CNS tumor was confirmed by patient health record and pathologist determination upon biopsy/resection. For reference genes within the tumor samples, the outputs of *FindAllMarkers* at the cluster level were used as input to CASSIA.

